# MEK inhibition enhances the antitumor effect of radiation therapy in *NF1*-deficient glioblastoma

**DOI:** 10.1101/2023.08.04.552061

**Authors:** Maria Ioannou, Kriti Lalwani, Abiola A. Ayanlaja, Viveka Chinnasamy, Christine A. Pratilas, Karisa C. Schreck

**Affiliations:** Department of Neurology, Johns Hopkins University School of Medicine, Baltimore, MD; Department of Oncology, Sidney Kimmel Comprehensive Cancer Center at Johns Hopkins University School of Medicine, Baltimore, MD; Department of Pediatric Oncology, Johns Hopkins University School of Medicine, Baltimore, MD

**Author notes:** Corresponding author: Karisa C. Schreck, Full name: Karisa C. Schreck, Mailing address: 201 N. Broadway, Rm. #9179, Baltimore, MD. **Additional information:**.

**Keywords:** glioblastoma, *NF1*, irradiation, MEK inhibition, homologous recombination, DNA damage repair

## Abstract

Individuals with neurofibromatosis type 1 (NF-1), an autosomal dominant neurogenetic and tumor predisposition syndrome, are susceptible to developing low-grade glioma (LGG) and, less commonly, high-grade glioma (HGG). These gliomas exhibit loss of the neurofibromin gene (*NF1)*, and 10-15% of sporadic HGG have somatic *NF1* alterations. Loss of NF1 leads to hyperactive RAS signaling, creating opportunity given the established efficacy of MEK inhibitors (MEKi) in plexiform neurofibromas and some individuals with LGG. We observed that *NF1*-deficient glioblastoma neurospheres were sensitive to the combination of a MEKi (mirdametinib) with irradiation, as evidenced by synergistic inhibition of cell growth, colony formation, and increased cell death. In contrast, *NF1*-intact neurospheres were not sensitive to the combination, despite complete ERK pathway inhibition. No neurosphere lines exhibited enhanced sensitivity to temozolomide combined with mirdametinib. Mirdametinib decreased transcription of homologous recombination genes and RAD51 foci, associated with DNA damage repair, in sensitive models. Heterotopic xenograft models displayed synergistic growth inhibition to mirdametinib combined with irradiation in *NF1*-deficient glioma xenografts, but not those with intact *NF1*. In sensitive models, benefits were observed at least three weeks beyond the completion of treatment, including sustained phospho-ERK inhibition on immunoblot and decreased Ki-67 expression. These observations demonstrate synergistic activity between mirdametinib and irradiation in *NF1*-deficient glioma models and may have clinical implications for patients with gliomas that harbor germline or somatic *NF1* alterations.

## Introduction

The mainstay of therapy for glioblastoma in adults for almost 20 years has been maximal safe surgical resection followed by radiation with concurrent oral alkylating chemotherapy, temozolomide (TMZ), but this regimen is modestly effective at best(1). In the 40% of patients whose tumors are deficient for O6-methylguanine-DNA methyltransferase (MGMT promoter is methylated), median survival improves to approximately 23 months; however in patients whose tumors are MGMT unmethylated, median survival remains less than 18 months^2,^(3). Detailed genomic characterization of glioblastoma has led to a deeper understanding of prognostic and predictive alterations, reflected in the most recent WHO Brain Tumor Classification(4). Despite these breakthroughs, our ability to exploit oncogenic genomic alterations remains limited, with many negative clinical trials over that time.(5)

ERK signaling has proven to be critical for glioblastoma cellular growth, as approximately 80% of all glioblastomas have genomic alterations that dysregulate this pathway, either through oncogenic activating mutations (e.g., EGFRvIII, BRAF^V600E^), or through loss of tumor suppressors (*PTEN, TP53, NF1*). Despite promising preclinical data, clinical trials of targeted therapies against EGFR, mTOR, PI3K and other commonly altered targets have been unsuccessful in improving overall survival in patients with glioblastoma (6),(7). Even with demonstrable target inhibition in human subjects, development has been limited by both suboptimal antitumor efficacy, intratumoral heterogeneity, and toxicity (8). Therapies with broader inhibition of ERK signaling, good brain penetration, and a tolerable toxicity profile are needed for evaluation in glioblastoma.

One approach is to identify therapeutic agents that synergize with current FDA-approved therapies (radiation or TMZ) potentially by augmenting DNA damage or preventing repair. DNA damage induced by radiation can be repaired via non-homologous end-joining (NHEJ) pathway or homologous recombination (HR), which is primarily involved in the repair of secondary double strand breaks (DSBs) that occur during S-phase following radiation, when the replication fork collapses at unresolved single-stranded DNA (ssDNA) lesions(9). Both NHEJ and HR are executed through multistep reactions essential for repairing the DNA damage successfully. ERK signaling is critical for maintaining DNA damage repair pathways in many cancers (10),(11),(12),(13). Specifically, these pathways regulate RAD51 expression, along with other proteins essential to HR, such as BRCA1 and BRCA2. Inhibition of ERK signaling with a MEK inhibitor suppresses the transcription of HR pathway genes in melanoma cell lines and renders cells susceptible to therapies that induce DSBs.(10)

To assess the role of ERK signaling on HR activity in glioblastoma we evaluate the effect of combined MEK inhibition (using the MEK inhibitor, mirdametinib) with cytotoxic therapy (radiation or TMZ) in a panel of glioblastoma neurospheres with genomic alterations conferring ERK dependence, specifically *NF1* loss or BRAF^V600E^ mutations, versus those without ERK dependence. Our findings demonstrate cooperative activity between MEKi and radiation in glioblastoma models, both *in vitro* and *in vivo*, with potential implications for human clinical trials.

## Materials and Methods

### Cell Culture Conditions

Human-derived glioblastoma neurosphere lines (HSR-GBM1, JHH-520, JHU-0879, JHU-1016B, JHH-136) with diverse *NF1* status were obtained and cultured in serum-free neurosphere media as previously described.(14) Adherent high-grade glioma lines with BRAF^V600E^ alteration and varied *NF1* status (B76, NGT-41, NMCG-1, DBTRG) were obtained and grown in attached culture media with 10-15% serum as previously described(15). LN229 was obtained from ATCC, T98G was a gift from the Laterra laboratory, and both were cultured as recommended. All cell lines used in our study were verified by short-tandem repeat profiling for cell line authentication at Johns Hopkins University Core Facility, tested regularly for *Mycoplasma* contamination, and passaged *in vitro* for fewer than three months after resuscitation.

### Cell Growth Assays

Cell lines were infected with NucLight red lentivirus and stably selected in puromycin. Successful insertion of the nuclear-restricted RFP was confirmed via fluorescence microscopy of stably infected cells. Cell growth was evaluated using an IncuCyte S3 live Imager by monitoring area of red nuclei every four hours, or the Cell Counting kit-8 (Dojindo) and read using a microplate spectrophotometer (Epotech) as described. For all growth assays, 1.5-3 × 10^3^ cells/well were plated in 96-wells plates and treated 24 hours after plating with the described conditions. Relative survival in the presence of drug was normalized to untreated controls after background subtraction(16). Synergy was calculated using the web-based SynergyFinder to analyze drug combination dose–response matrix data. To determine whether the drugs showed a combinatorial or additive effect, a Synergy Score was calculated to characterize the strength of synergistic interaction(17),(18). For all experiments at least three independent replicates were performed.

Soft-agar assay was performed as described previously(19). Briefly, 100 to 150 × 10^3^ cells were mixed with 1% agar treated with either DMSO or mirdametinib (10 or 30 nM) and plated over a base layer in 6-well plates. Plates were radiated as described and incubated at 37 °C for three weeks with media changes and drug repletion one to two times weekly. Colonies were stained with crystal violet for one hour and imaged via ChemiDoc Touch Imaging System (Bio-Rad). The measurements were based on three replicates for each condition and each experiment was repeated at least three times. Images captured within a single experiment were taken at the same magnification and exposure time. Data were analyzed using ImageJ.

### Immunoblotting

Cells were prepared and processed as previously described(20). Briefly, cells were disrupted on ice in NP40 lysis buffer. Protein concentration was determined with Pierce BCA protein assay kit (#23227, Thermo Fisher Scientific). Equal amounts of protein were separated by SDS-PAGE, transferred to nitrocellulose membranes, immunoblotted with specific primary and secondary antibodies, and detected by chemiluminescence with the ECL detection reagents, Immobilon Western chemiluminescent HRP substrate (#WBKLS0500, Millipore), or Pierce ECL Western blotting substrate (#32106, Thermo Fisher Scientific). The membranes were imaged using ChemiDoc touch imaging system (Bio-Rad). Representative blots of three to four independent experiments are shown. Antibodies against total ERK, phospho-ERKthr202/tyr204, phospho-MEKser217/221, cleaved PARP, RAD51, BRCA2 and GAPDH were obtained from Cell Signaling Technology. NF1 (A300-140A) was obtained from Bethyl. MGMT (sc-166528) was obtained from Santa Cruz Biotechnology.

### Real-time PCR

RNA was extracted from treated cells and reverse transcription performed as previously described(19). qPCR was performed using the Promega 2-step kit on a CFX96 Real-Time PCR System (Bio-Rad) according to the manufacturer’s instructions. Three independent experiments (with biological triplicate) were completed for each condition using previously published primers for RAD51, BRCA2 and GAPDH(21). Values were normalized to the housekeeping gene GAPDH using the ΔΔCT method (22).

### Flow Cytometry

Cells were plated in 6-well plates, irradiated the same day and treated immediately afterwards with mirdametinib. Cells were collected after 72 hours and fixed with 70% ethanol. They were stained with Propidium Iodine (PI/RNase staining-Cell Signaling Technologies), then sorted using a FACS Celeste machine. For Annexin V staining, cells were harvested and resuspended in PBS, then treated with Annexin V, Alexa Fluor 488 conjugate (Invitrogen 2413465) and Propidium Iodide prior to flow cytometry. Results were analyzed using the FlowJo software.

### Immunofluorescence

Cells were grown in chamber slides (Lab Tek II, Chamber Slide cat. No. 154534) before fixation (4% paraformaldehyde in PBS for 15 minutes) and permeabilization (0,2% Triton X-100 in PBS). Cells were blocked with goat serum for 2 hours at room temperature and incubated with RAD51(1:500 no. ab133534 Abcam) or yH2AX (1:200 no. 05-636, Millipore) antibody overnight at 4 °C and then with secondary antibody (1:1000 goat anti-mouse Alexa Fluor 488 or 1:2000 goat anti rabbit Alexa Fluor 647) and DAPI (Thermo Fisher). Image acquisition was performed with a confocal microscope (LEICA SP8 Confocal Microscope).

### Immunohistochemistry

Immunohistochemistry (IHC) was performed at Johns Hopkins IHC core facility. IHC for Ki-67 and pERK was performed on formalin-fixed, paraffin-embedded sections on a Ventana Discovery Ultra autostainer (Roche Diagnostics) using a protocol described previously(23). Ki-67 quantification and image analysis was performed with Qupath to score Ki-67 in tumor tissue for a cohort of nine slices per condition.

### Radiation therapy

Cells were plated in 96 well plates and irradiated with 0–10 Gy using a GammaCell irradiator with a ^137^Cs source at 50 cGy/min. For *in vivo* experiments, mice were treated using the Small Animal Radiation Research Platform (SARRP), which was able to target the tumor tissue as appropriate(24). The tumors were irradiated with a circular beam of 1-cm diameter with three consecutive daily fractions of 2 Gy each.

### Animal Experiments and Ethics Statement

NOD scid gamma (NSG) female and male mice were purchased from Johns Hopkins University Animal Resources Core. All mouse experiments were approved by the Institutional Animal Care and Use Committee at Johns Hopkins under protocol # MO21M387. Glioblastoma neurosphere lines were implanted subcutaneously in 6 to 8 week-old NSG mice as described.(25) Drug treatment was started when tumor size reached roughly 50-200 mm^3^, at which time mice were randomized into treatment groups and treated accordingly. Mirdametinib (1.5 mg/kg, dissolved in 0.5% hydroxypropyl methyl cellulose and 0.2% Tween 80) was administrated by oral gavage twice daily, based on mean group body weight, with treatment schedule of 7 days on/0 days off. Irradiation was administered in 3 daily fractions of 2 Gy. Tumors were measured twice weekly by caliper in two dimensions, and tumor volume was calculated by: W x L^2^ where L is the longest diameter and W is the width. Mice were treated for 28 days and monitored closely for toxicity and signs of tumor growth or neurological impairment. Tumors were extracted, photographed, and fixed in formalin.

### Data analysis and Statistical Methods

Graphing, IC50 calculations, and statistical analysis were performed using GraphPad Prism, version 7. ANOVA or multiple t tests were used to evaluate difference between conditions.

### Data Availability

The data generated by the authors are available upon request to the corresponding author.

## Results

### MEK inhibition suppresses HR pathway activity in NF1-deficient glioblastoma

We started by determining ERK-dependence in our cell line models based on the presence of a genomic alterations in the ERK signaling cascade (*NF1* loss or *BRAF* alteration) as well as dose-dependent growth inhibition to the small molecule allosteric MEK1/2 inhibitor, mirdametinib (26) (also known as PD0325901, MEKi), at physiologically relevant doses. We compared existing genomic alterations for all lines and to *NF1* expression status determined by immunoblot **(Fig. 1A)** Based on integrated analysis, we identified two lines as *NF1* intact (HSR-GBM1, JHU-0879), three as *NF1* deficient (JHH-136, JHH-520, JHH-1016B), three with BRAF^V600E^ and *NF1* intact (B76, DBTRG, NMCG1) and one with BRAF^V600E^ and *NF1* deficiency (NGT41). Serum-free neurosphere and adherent glioblastoma cell lines were all treated with a dose range of MEKi. Glioblastoma models exhibited varied sensitivity to MEKi, with highest sensitivity in those with loss of *NF1* or BRAF^V600E^ alterations—consistent with an ERK-dependent phenotype—and little growth inhibition in those with intact *NF1* and BRAF, despite adequate inhibition of ERK activity (phospho-ERK) in all cell lines, as previously reported (27)**(Fig. 1B-D)**.

**Figure 1:**
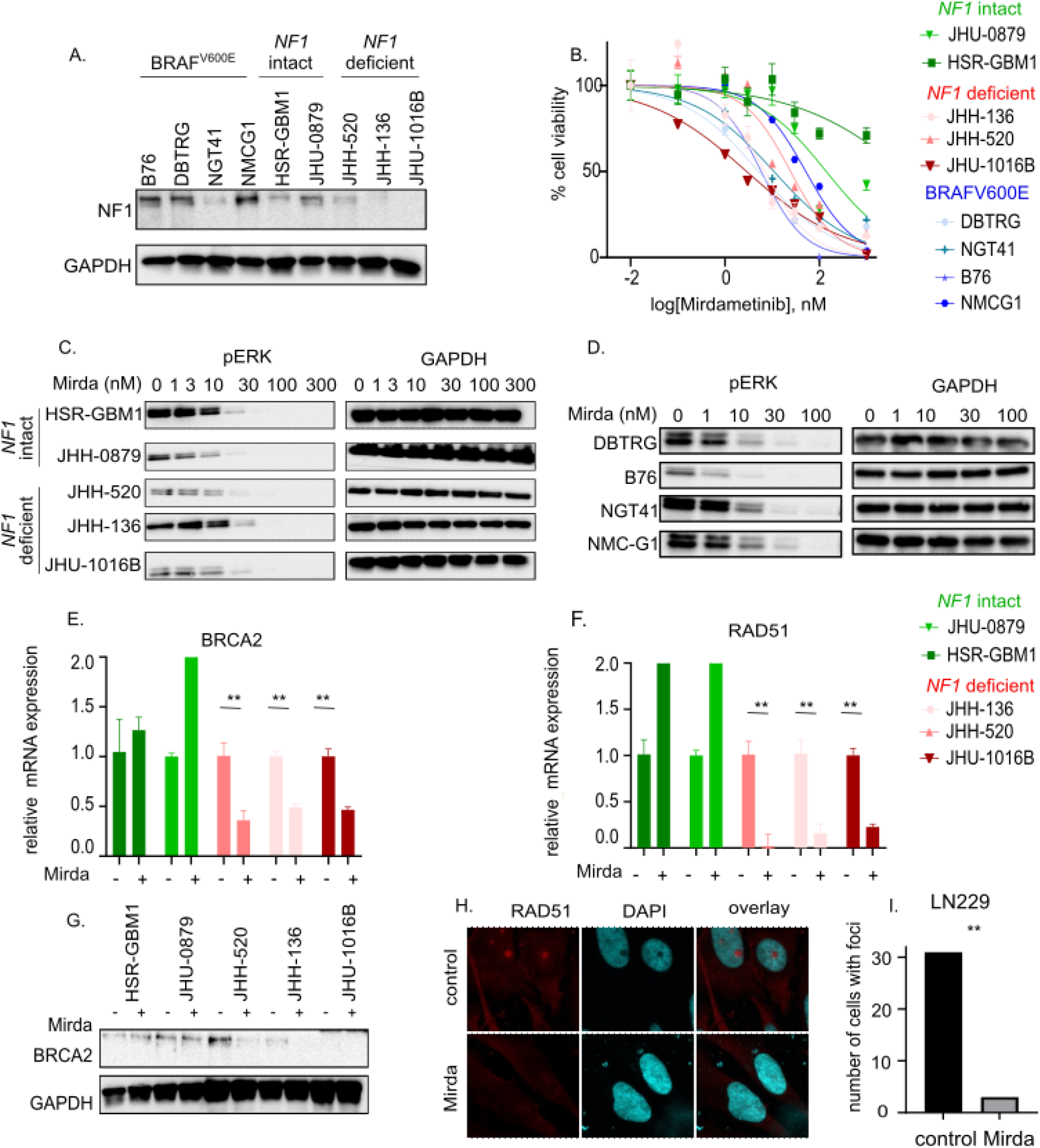
*NF1* alteration determines decreased HR expression following MEK inhibition. **A)** *NF1* status was verified using immunoblot in four adherent glioma lines and five neurosphere models. **B)** Cell lines were exposed to increasing concentrations of the MEK inhibitor mirdametinib for 96 hours and cell viability was evaluated using the CCK-8 cell viability assay, or **C/D)** treated with DMSO or varied doses of mirdametinib for one hour and pERK levels were assessed using immunoblot. **E/F)** Five neurosphere lines, two *NF1* intact (green), and three *NF1* deficient (red) were treated with DMSO or 30 nM of mirdametinib for 24 hours and relative mRNA expression of *BRCA2* and *RAD51* was assessed by qPCR (ddCt). **G)** BRCA2 immunoblot after 24 hours of treatment with 30 nM of mirdametinib. **H/I)** The *NF1-/-* adherent glioma line LN229 was treated with 30 nM of mirdametinib for 24 hours, fixed and stained for DAPI and RAD51. Image acquisition and quantification of RAD51 foci was performed using the LEICA confocal microscope and software. All panels show representative data from at least three independent replicates. **p<0.005 by student t-test

Given prior reports of ERK-dependent regulation of homologous recombination (HR)(10,28,29), we sought to evaluate the putative relationship in glioblastoma. We quantified expression of *BRCA2* and *RAD51* mRNA, two genes critical for HR, in cells treated with DMSO or MEKi. We observed decreased transcription of both genes following 24 hours of treatment with MEKi in neurosphere lines with *NF1* deficiency, though not in those with intact *NF1* (**Fig 1E-F**). We observed decreased BRCA2 protein expression (**Fig. 1G**), and fewer RAD51 nuclear foci following MEKi therapy as compared with controls in an *NF1*-deficient adherent glioblastoma line, consistent with ERK-dependent regulation of HRD **(Fig 1H, I)**.

### MEK inhibition sensitizes glioblastoma with loss of *NF1* to radiation

We next evaluated whether the MEKi, mirdametinib, would sensitize glioblastomas to standard treatments by decreasing DNA repair. The effect of a single fraction of radiation on cell growth was analyzed and we observed dose-dependent inhibition and cell death in all cell lines. We identified 5 Gy as the dose with modest biological effect but incomplete cell killing across all neurosphere lines **(Suppl. Fig. 1A, B)**, akin to the state in human patients with glioblastoma treated with radiotherapy in whom cancer recurs over time. We additionally evaluated sensitivity to temozolomide (TMZ) monotherapy in our lines **(Suppl. Fig. 1C)**, which in glioblastoma is determined by MGMT expression(30). As expected, lack of MGMT expression (HSR-GBM1, JHH-520, JHU-1016B) conveyed sensitivity to TMZ, while intact MGMT expression (JHU-0879 and JHH-136; **Suppl. Fig. 1D**) was associated with resistance.

We assessed the effect of combining MEKi therapy with irradiation. Across all *NF1*-deficient neurospheres, but not *NF1*-intact lines, we observed decreased cell growth and sphere formation in response to combined therapy **(Fig. 2A, C)**. Adherent cell lines with BRAF^V600E^ or *NF1*-deficiency also displayed additive or synergistic response to MEKi combined with radiation **(Fig. 2B, D)**. Additive effects were not observed for *NF1*-intact neurospheres or adherent cell lines. We evaluated the effect of combined therapy on the clonogenic potential of glioblastoma stem cells using a soft agar colony-forming assay and observed decreased sphere formation with combined treatment compared to either monotherapy in *NF1*-deficient but not *NF1*-intact neurospheres **(Fig. 2E, F)**.

**Figure 2:**
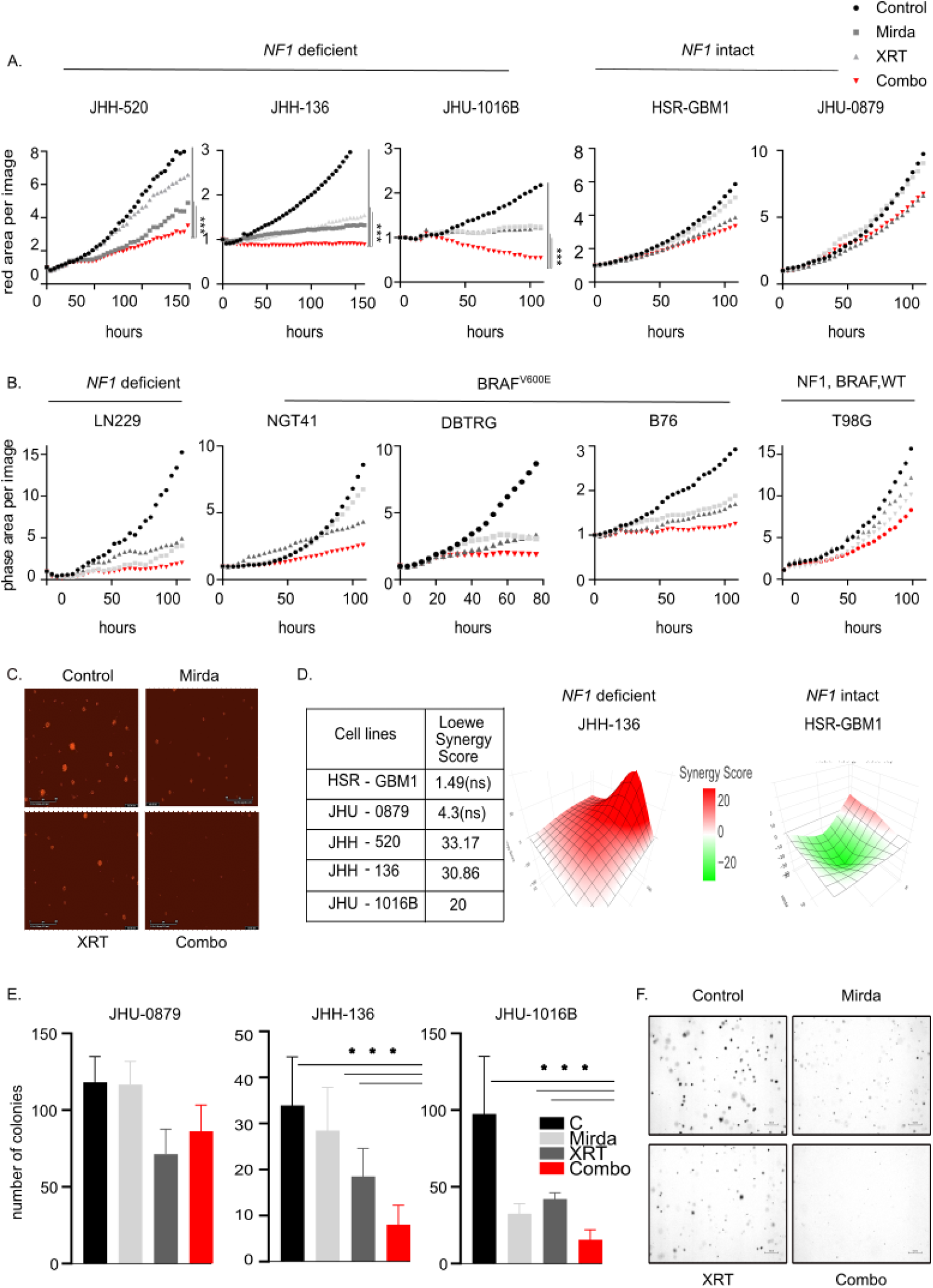
The combination of radiation and mirdametinib synergistically inhibits cell growth in *NF1* deficient glioma. **A/B)** Neurospheres expressing a red nuclear label or adherent glioma lines were treated with DMSO, 5 Gy of radiation, 10 nM of mirdametinib or their combination for up to seven days. The red area per image or the phase confluence was monitored by the Incucyte real-time system, normalizing to corresponding 0-hour scan, for each frame. **C)** Representative images for the *NF1*-deficient neurosphere line JHH-136 after 72 hours of treatment. **D)** Synergy scores were calculated using the Loewe model. **E/F)** One *NF1*-intact (JHU-0879) and two *NF1-*deficient lines (JHH-136, JHU-1016B) were embedded in soft agar, treated with DMSO, 10 nM of mirdametinib, 5 Gy of radiation or their combination for three weeks and colonies were stained with crystal violet to quantify colony forming ability. Representative images for JHU-1016b are shown after two weeks of treatment. All panels show representative data from at least three independent replicates. ***p<0.01 by one-way ANOVA or by student t-test

To further characterize the effect of combined treatment, we investigated the effect on cell cycle progression. In *NF1*-deficient neurospheres, the combination of MEKi and radiation produced a higher sub-G1 population and increased cell death, along with decreased populations in S and G2 phases **(Fig. 3A)**. This effect was not observed in *NF1*-intact neurospheres treated with the combination. *NF1-*deficient neurospheres also demonstrated increased apoptosis in response to combined therapy as compared with either monotherapy when measured by Annexin V **(Fig. 3B, C)**.

**Figure 3:**
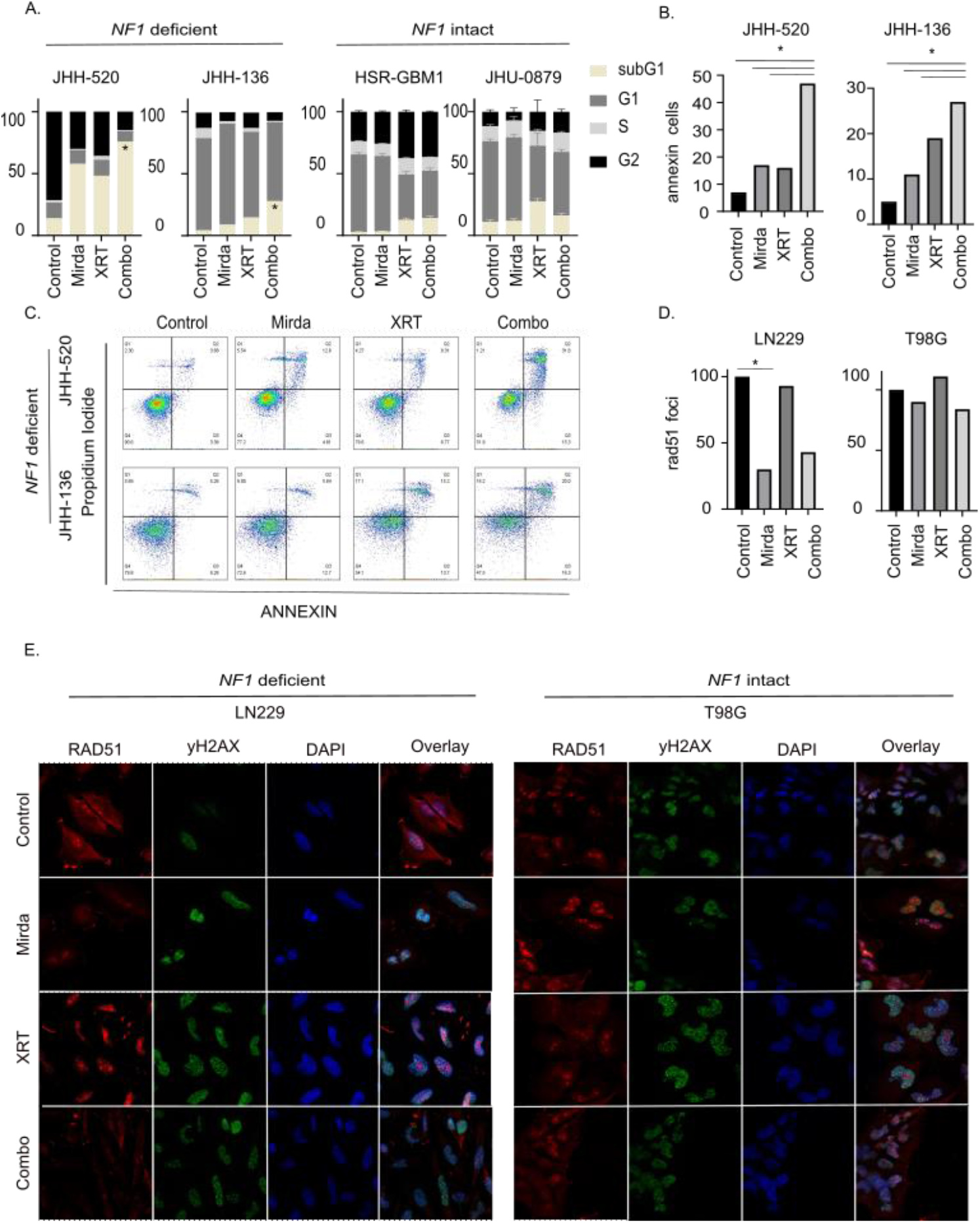
Combination of radiation and mirdametinib inhibits cell cycle and induces apoptosis in *NF1* deficient glioma. **A)** Two *NF1* deficient (JHH-520, JHH-136) and two *NF1* intact (HSR-GBM1, JHU-0879) neurosphere lines were treated with DMSO, 10 nM of mirdametinib, 5 Gy of radiation or their combination for 72 hours. Cells were fixed with 70% ethanol and stained with propidium iodide/RNase staining solution, and then analyzed by flow cytometry. **B/C)** JHH-136 and JHH-520 cells were treated as described for 72 hours, stained with annexin V, and analyzed by flow cytometry. The percentage of annexin positive cells was calculated from three independent experiments and is shown in bar plots. **D/E)** *NF1* deficient (LN229) and *NF1* intact (T98G) cell lines were treated with DMSO, 10 nM mirdametinib, 5 Gy of radiation or the combination, fixed with 4% paraformaldehyde, washed with PBS, permeabilized and incubated with DAPI, yH2AX or RAD51. Images were acquired and analyzed using the LEICA software. All panels show representative data from at least three independent replicates. *p<0.05 by student t-test.

We evaluated the effect of combined therapy on HR protein expression. The number of RAD51 foci decreased with MEKi treatment in *NF1*-deficient cells, irrespective of whether radiation had occurred, suggesting decreased capacity for DNA damage repair using the HR pathway. In both *NF1*-intact and deficient lines, radiation resulted in increased DNA double-strand damage as quantified by γH2AX **(Fig. 3D, E)**.

### Effectiveness of radiation and MEKi in *in vivo* models of *NF1-*deficient glioblastoma

Given the synergistic combination of MEKi, mirdametinib, with irradiation *in vitro*, we evaluated the effect of combined therapy in heterotopic xenograft models. We observed a transient increase in the volume of tumors treated with radiation monotherapy that subsided within three weeks, consistent with transient inflammation. Overall, *NF1*-deficient heterotopic xenografts (JHH-520 and JHH-136) exhibited decreased xenograft growth with combined treatment as compared to monotherapy **(Fig. 4A-C)**. This was true whether the MEKi was administered simultaneously with irradiation or following the completion of radiation, but the benefit was most durable in mice treated concurrently, persisting for at least four weeks following the completion of treatment **(Suppl. Fig 2A)**. The combination failed to demonstrate a benefit in *NF1*-intact xenografts **(Fig. 4B)**. Immunoblot from tumors obtained following the last treatment confirmed ongoing suppression of ERK phosphorylation in arms with MEKi for *NF1* deficient tumors **(Fig. 4D, Suppl. Fig. 2B)**, demonstrating that phospho-ERK suppression alone is insufficient to inhibit tumor growth. Histopathologic analysis of tumor tissue from individual mice collected 28 days after the start of treatment confirmed a significant decrease in proliferation (Ki-67) in the combination arm, and decreased p-ERK in both conditions treated with MEKi **(Fig. 4E, F)**. Together, these findings support the observation that MEKi potentiates the effect of radiation in *NF1-*deficient glioblastoma, improving the amplitude and durability of treatment.

**Figure 4:**
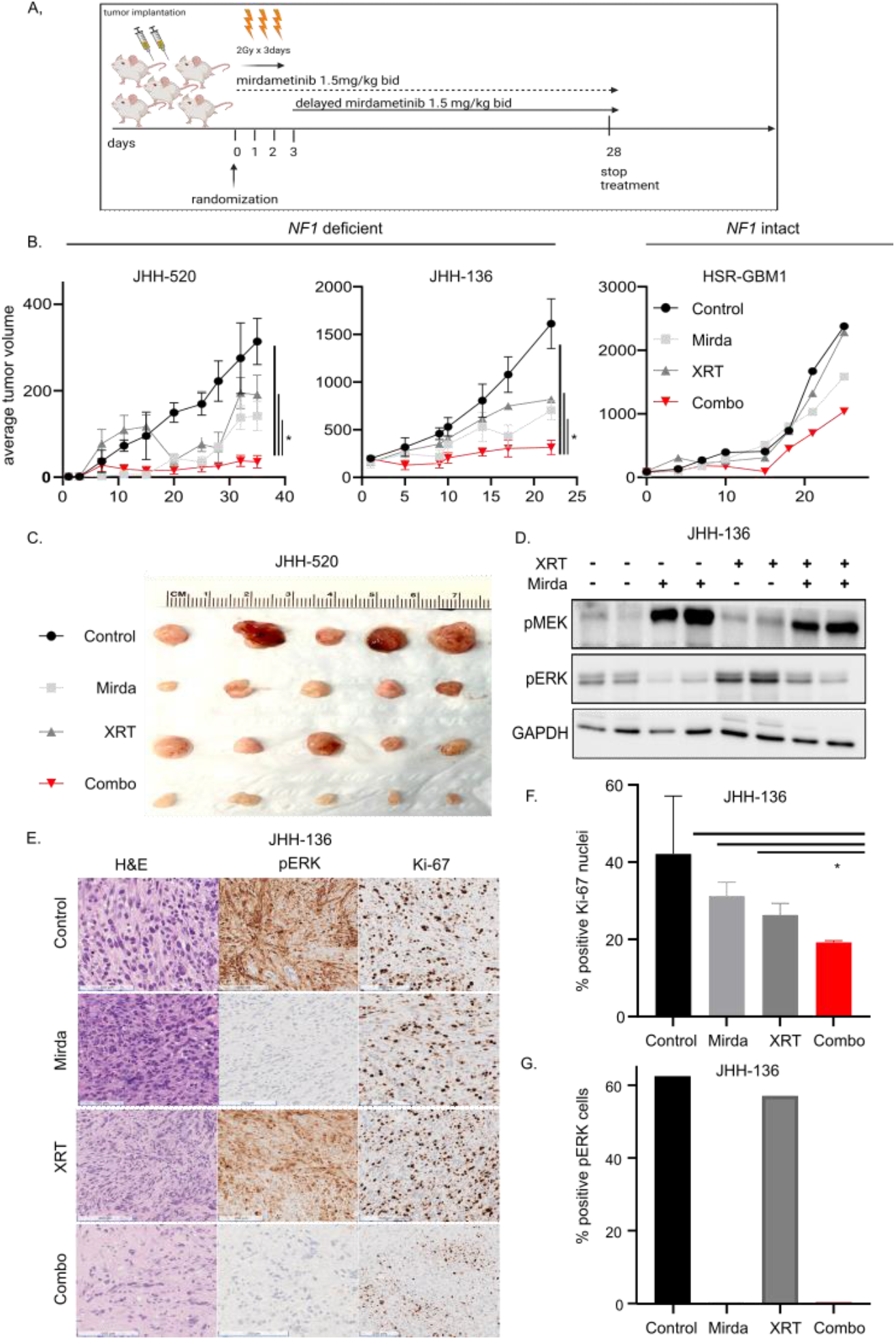
MEKi plus radiation is active against *NF1* deficient glioma models *in vivo*. **A/B)** Athymic NSG mice implanted with *NF1* deficient (JHH-520, JHH-136) or *NF1* intact (HSR-GBM1) cells were treated with vehicle, mirdametinib (1.5 mg/kg) via oral gavage twice daily for 4 weeks, radiation (fractionated 2 Gy in 3 days) or the combination. Tumor size was measured twice weekly. Average tumor was plotted over the time course of treatment days (n=8-10 tumors per arm). **C)** Five tumors from each cohort of JHH-520 were collected after completing 4 weeks of treatment with representative images shown. **D)** Two tumors from each arm of JHH-136 were collected five hours after the last treatment dose, lysed and subjected to immunoblotting with indicating antibodies, or **E/F/G)** fixed in 10% NBF and stained for Ki-67 or p-ERK with representative images shown. *p<0.05 by student t-test.

### MEKi does not potentiate temozolomide efficacy in glioblastoma models

Given its role as standard therapy for glioblastoma, we evaluated the combination of TMZ with MEKi. TMZ functions by adding methyl groups (alkylation) to guanine or adenine, causing missense substitutions during DNA replication and creating mismatched base pairs (31). This effect triggers DNA damage repair through mismatch repair and base excision repair mechanisms, rather than through homologous recombination. We hypothesized that there would be no synergy between TMZ and MEKi given DNA repair from TMZ functions independently of HR. We evaluated the combination of TMZ with MEKi but did not observe any synergistic effect on cell growth inhibition or cell death in any neurosphere lines **(Fig. 5A, B)**. Furthermore, histopathologic analysis of tumor tissue from individual mice collected 28 days after the start of treatment showed no difference in proliferation (Ki-67; **Suppl. Fig. 2D)**.

**Figure 5:**
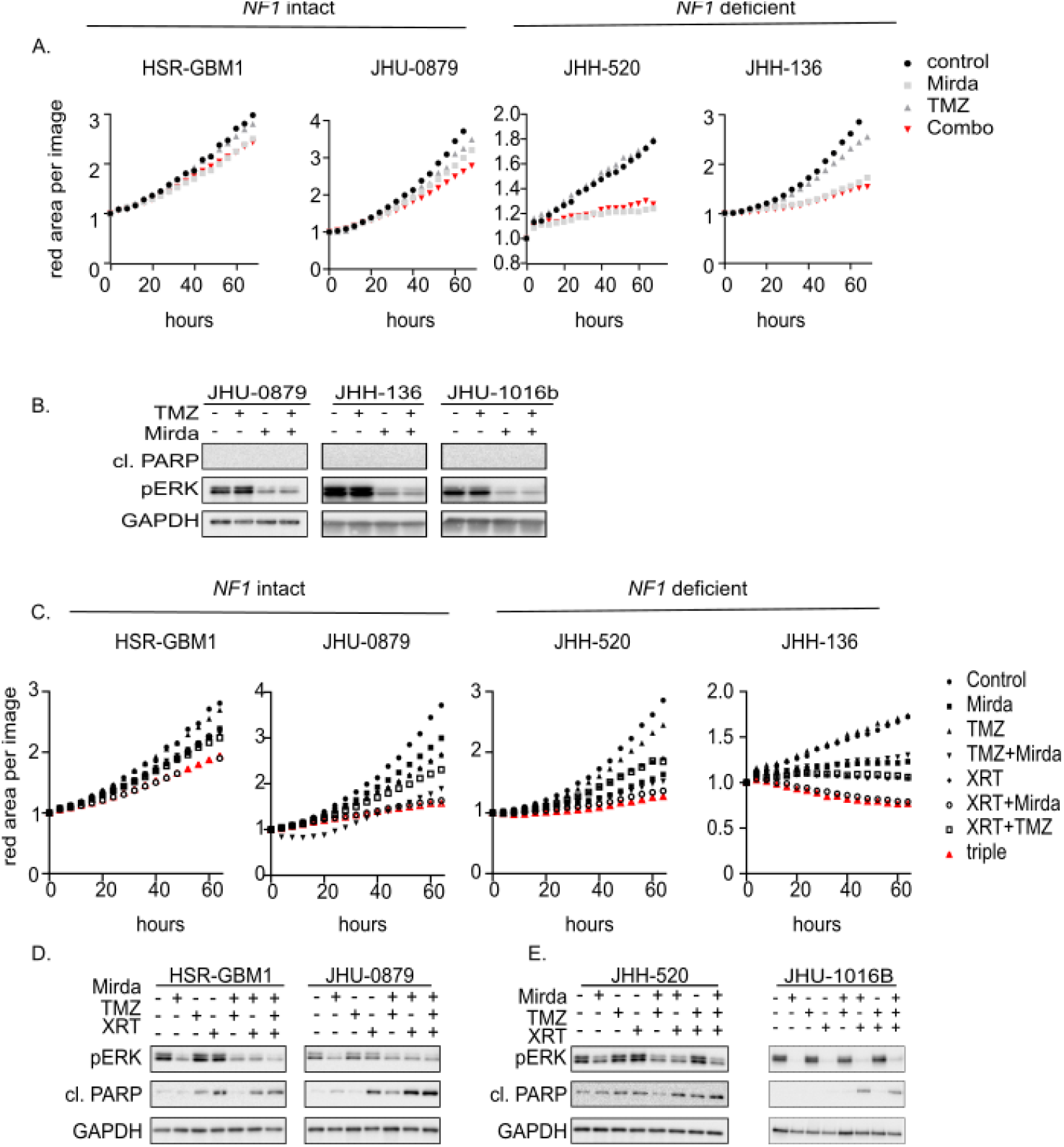
TMZ does not provide any additional benefit to the combination of MEKi and radiation. **A)** Neurospheres were treated with DMSO, 10 uM TMZ, 10 nM mirdametinib or their combination for 72 hours and cell growth was assessed using the Incucyte real time system or **B)** lysed and proteins were quantified using the listed antibodies. **C)** Two *NF1* deficient and two *NF1* intact neurosphere lines were treated with DMSO, TMZ 10uM, mirdametinib 10nM, 5 Gy, radiation, or their combinations for 72 hours. The red area per image was monitored by Incucyte real-time imaging system and normalized to corresponding 0-hour scan. **D/E)** Cells were treated as C and the indicating proteins were assessed using immunoblot.

Lastly, we evaluated the addition of MEKi to radiation and TMZ, the standard first-line therapy for glioblastoma. We observed a combinatorial effect of radiation with MEKi, but no enhanced growth inhibition or increased cell death with the addition of temozolomide **(Fig. 5C, D)**. This observation extended to heterotopic xenograft models, where MEKi with radiation was equivalent or superior to every other pairwise combination **(Suppl. Fig. 2B)**. Interestingly in *NF1*^*-/-*^ cell lines, apoptotic markers such as cleaved PARP demonstrated more induction for the combination of radiation with MEKi over any other combination, including the addition of TMZ or TMZ with radiation alone **(Fig. 5E)**.

## Discussion

Radiation therapy is a cornerstone of glioblastoma treatment and plays a critical role in controlling tumor growth. Combined with oral TMZ chemotherapy, it comprises first line therapy for glioblastoma and other high-grade gliomas in adults. Unfortunately, glioblastoma cells can overcome the effects of standard treatment, which is focused on inflicting catastrophic DNA damage, through aberrant DNA repair mechanisms and cell-cycle regulation. Here, we demonstrate that *NF1*-deficient glioblastoma lines are ERK-dependent, with partial growth sensitivity to MEKi monotherapy. They also exhibit ERK-dependent expression of the homologous recombination DNA repair pathway, where MEKi creates a transient homologous recombination deficiency (HRD)-like state in NF1-deficient cells. This vulnerability is exploitable by treating with MEKi and radiotherapy, effecting enhanced growth inhibition and cell death both *in vitro* and *in vivo*. These findings have potential clinical implications for sporadic glioblastoma with *NF1* loss, as well as for patients with high-grade glioma in the background of neurofibromatosis type 1.

Genomic alterations conferring HRD are well known to confer increased radiosensitivity. Mutations in *BRCA1* or *BRCA2* represent examples of classic cancer-associated DNA repair defects, conferring therapeutic susceptibility to agents that stall replication forks and induce double-strand DNA breaks(32). These breaks are typically repaired via the NHEJ or HR pathways(33). Moreover, alterations in the isocitrate dehydrogenase gene (IDH1/2) occur in glioma, conferring an HR deficient state and increased sensitivity to radiotherapy in patients(34). Here, we demonstrate that *NF1*-deficient glioblastomas exhibit HRD due to decreased transcription following treatment with MEKi, which in turn confers enhanced sensitivity to irradiation. The overall effect is accumulation of unrepaired DNA damage leading to apoptosis. The dependence of HR expression on ERK signaling appears to be unique to glioblastoma with loss of *NF1* or BRAF^V600E^ alterations, suggesting an ERK-dependent phenotype. This is consistent with prior observations that melanoma and other cancers with RAS-dependence (due to *BRAF, NF1*, or *NRAS* alterations) demonstrate ERK-dependent HR, creating an exploitable vulnerability (10,12,29).

Our preclinical experimentation was conducted using a combination of *NF1*-deficient and *NF1*-intact glioblastoma models derived from sporadic glioblastoma in adult patients. Low-grade glioma, particularly optic pathway glioma, is quite common in patients with neurofibromatosis type 1 (NF1, heterozygous germline loss of neurofibromin type 1). LGG in patients with NF1 respond to MEKi alone, and even the occasional HGG may respond as well (35–37). However, there is no standard treatment for HGG in patients with NF1 and radiation is often incorporated into therapy. In patients with sporadic glioblastoma, on the other hand, mono or bi-allelic alterations in *NF1* lead to loss of protein expression or function in ∼15%. We expect the findings of this study, conducted in sporadic glioblastoma models, are also relevant for HGG from patients with NF1, but this hypothesis will need to be confirmed in future studies.

Attempts to combine radiation with molecularly targeted therapies in the past have been ineffective in glioblastoma. There have been several clinical trials evaluating the combination of EGFR inhibitors and radiation, but most of them were phase I studies focused on dose-finding and safety rather than efficacy. Importantly, in none of the studies did a significant safety signal emerge, indicating that the addition of a small molecule inhibitor to radiation was not more dangerous for patients (38),(39),(40). One significant obstacle has been the blood-brain barrier, which is impervious to most chemotherapies and small molecule inhibitors, including most EGFR inhibitors. MEK inhibitors, conversely, have proven blood-brain barrier penetration as evidenced by significant tumor responses pediatric low-grade glioma(35,37,41), resulting in FDA-approval of two different MEKi, either alone or in combination with a BRAF inhibitor. Mirdametinib, a brain-penetrant, high selective allosteric MEK1/2 inhibitor, is currently undergoing clinical trial evaluation in patients with pediatric LGG (NCT04923126) (42,43).

We did not observe any enhanced efficacy for the combination of TMZ with MEKi. This limitation is likely due to the fact that TMZ induces the formation of mismatched base pairs, which are detected and processed by mismatch repair (MMR) enzymes (44), rather than HR. Because this is a different repair mechanism, the addition of the MEKi mirdametinib to TMZ did not demonstrate any synergy in our models. These findings are in agreement with groups that have previously evaluated this combination, showing no further benefit to the addition of MEKi to TMZ (45,46).

In conclusion, we have demonstrated that MEKi blocks HR in *NF1*-deficient glioblastoma resulting in synergy with radiation in a cohort of glioblastoma neurosphere lines and *in vivo* models. While confirmation in orthotopic xenografts is needed for further validation, these observations have potentially important implications for patients with NF-1 associated glioma, as well as for those with *NF1-*deficient sporadic glioblastoma.

## Authors’ contributions

Conception and design: K.C.S., C.A.P.

Development and methodology: M.I., K.C.S., C.A.P.

Acquisition, analysis, and interpretation of data: M.I., K.L., V.C., A.A., K.C.S.

Writing, review, and revision of manuscript: M.I., K.L., A.A., K.C.S., C.A.P.

## Acknowledgements

The authors acknowledge Drs. Charles Eberhart and Gregory Riggins for providing neurosphere lines and Esteban Velarde for providing technical support with irradiation. We are also grateful for scientific discussions with Drs. Lindy Zhang, Ava Wang and Patience Odeniyide.

