## Supplemental Figure 1 for "MEK inhibition enhances the antitumor effect of radiation therapy in *NF1*-deficient glioblastoma"

**SUPPL. FIG 1**

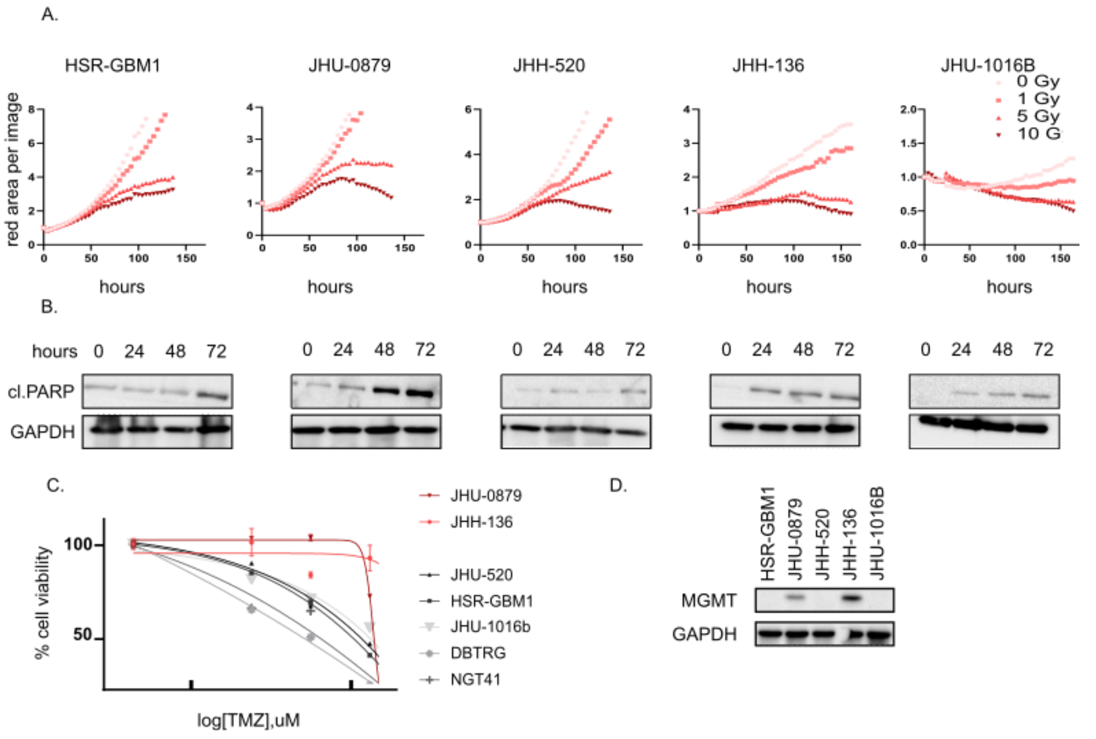


Supplementary figure 1: MGMT status determines TMZ sensitivity. **A)** Five neurosphere lines were treated with 0, 1, 5, or 10 Gy of radiation for up to seven days. Cell growth was monitored by the Incucyte real-time system, normalizing to corresponding 0-hour scan, for each frame or **B)** cells were lysed and proteins were quantified using the listed antibodies. **C)** Cell lines were exposed to increasing concentrations of TMZ for 96 hours and cell viability was evaluated using the CCK-8 cell viability assay. **D)** MGMT status, determining sensitivity to temozolomide, was determined by immunoblot.
