## Supplemental Figure 2 for "MEK inhibition enhances the antitumor effect of radiation therapy in *NF1*-deficient glioblastoma"

**SUPPL. FIG 2**

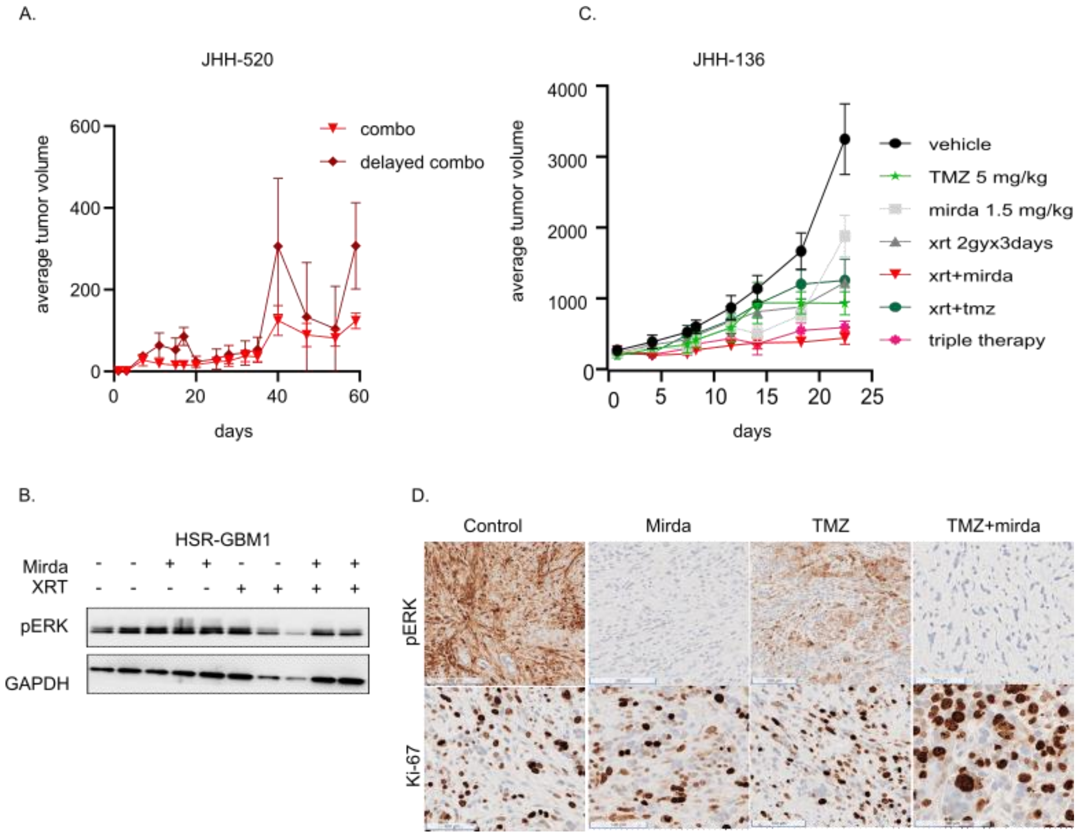


Supplementary figure 2: MEKi plus TMZ is not active against *NF1* deficient glioma models *in vivo*. **A)** Athymic NSG mice implanted with *NF1* deficient JHH-520 were treated with the combination of radiation (fractionated 2 Gy in 3 days) and mirdametinib (1.5 mg/kg) via oral gavage, starting simultaneously (combo) or after the completion of radiation (delayed combo). **B)** Xenografts from *NF1* intact HSR-GBM1 were lysed and indicating proteins were assessed by immunoblot. **C)** Athymic NSG mice implanted with *NF1* deficient JHH-136 were treated with vehicle, mirdametinib (1.5 mg/kg) via oral gavage twice daily for 4 weeks, radiation (fractionated 2 Gy in 3 days), TMZ (5 mg/kg) via oral gavage daily for 4 weeks or their combinations. **D)** Tumors from JHH-136 were fixed in 10% NBF and stained for Ki-67 or p-ERK with representative images shown
